# GeneCodis 4: Expanding the modular enrichment analysis to regulatory elements

**DOI:** 10.1101/2021.04.15.439962

**Authors:** A. García-Moreno, R. López-Domínguez, A. Ramirez-Mena, A. Pascual-Montano, E. Aparicio-Puerta, M. Hackenberg, P. Carmona-Saez

## Abstract

GeneCodis is a web-based tool for functional enrichment analysis that allows researchers to integrate different sources of annotations. It extracts sets of significant concurrent annotations and assigns a statistical score to evaluate those that are significantly enriched in the input list. Since its first release in 2007, it has been widely employed to analyze lists of genes in order to interpret the underlying biological mechanisms. Here we present GeneCodis 4, a new release that expands the functional analysis provided by the application, to accept regulatory elements, including lists of, transcription factors, CpG sites and miRNAs. It also incorporates new annotation databases and improved interactive visualizations functionalities to explore results. GeneCodis 4 is freely available at https://genecodis.genyo.es.

## INTRODUCTION

Functional enrichment analysis is a widely used method to discover biological annotations that are over-represented in a list of genes with respect to a reference list. These significant annotations are used to interpret the biological mechanisms and molecular pathways that are underlying the analyzed phenotype.

GeneCodis was one of the first applications that allowed researchers to integrate different sources of annotations to perform modular enrichment analysis (Carmona-Saez et al., 2007). It combines different types of annotations and extracts sets of terms that are jointly associated to sets of genes from the input list. Since this first release, the application has been continuously updated due to its extensive use and demand (Nogales-Cadenas et al., 2009; Tabas-Madrid et al., 2012).

In the last decade, new omics techniques have fuelled the extension of high-throughput analysis to other biological entities beyond genes or proteins. These methods are widely used for genome analysis, global methylation, miRNAs discovery and profiling or transcription factor (TF) analysis, among many other applications. The analysis of epigenetics patterns or regulatory elements have been an increasing focus of research in many contexts, such as development biology or characterization of molecular mechanisms in complex diseases (Schübeler, 2015). In this context, in a similar way than the analysis of genes or proteins, these experiments generate large lists of TFs, CpG sites or miRNAs and the challenge is to understand the key biological processes that are acting in such experiments.

Therefore, there is an increasing need for dedicated methods and tools for functional analysis of these types of experiments. In this context, different groups are proposing dedicated tools for functional analysis of miRNAs (Garcia-Moreno and Carmona-Saez, 2020) or methylation data (Ren and Kuan, 2019). Most of these are developed with the consideration of a well-known bias, in the case of using methylation data, several of the terms that seem enriched would also be identified with randomized data (Geeleher et al., 2013), which also happens with miRNAs (Bleazard et al., 2015), concretely, cancer and cell cycle terms are enriched (Godard and van Eyll, 2015).

Other methods aim to integrate multi omics data to derive biological knowledge, although they are not provided as tools but as pipelines to be used with specific datasets. ActivePathways (Paczkowska et al., 2020) is a recent functional analysis method that uses data fusion techniques to integrate different omics data. Complex methodologies like ActivePathways are not web-based unlike popular enrichment tools that are specialised in a single type of biological element. For example, modEnrichr (Kuleshov et al., 2019), GREAT (McLean et al., 2010) and miEEA (Kern et al., 2020) study the functional implications of lists of protein-coding genes, genomic regions and miRNAs, respectively.

In this work, the functionality of GeneCodis is extended to the analysis of regulatory elements. The web tool can now be used to the analysis of lists of TFs, CpG sites and miRNAs, besides to genes and proteins, providing all the functionalities that have been successfully applied to the analysis of genes lists to experiments that interrogate regulatory elements. In this new release, the application provides new visualization capabilities, extended the sources of annotations and supported databases and an updated and simplified back-end and frontend of the tool.

## MATERIAL AND METHODS

### Data Collection

#### Gene/protein and regulatory elements data collection

The new version of GeneCodis supports analysis for gene, protein, TFs, CpG sites and miRNAs identifiers from 16 different species, including main model organisms, that are listed in Figure 1.

**Figure 1.**
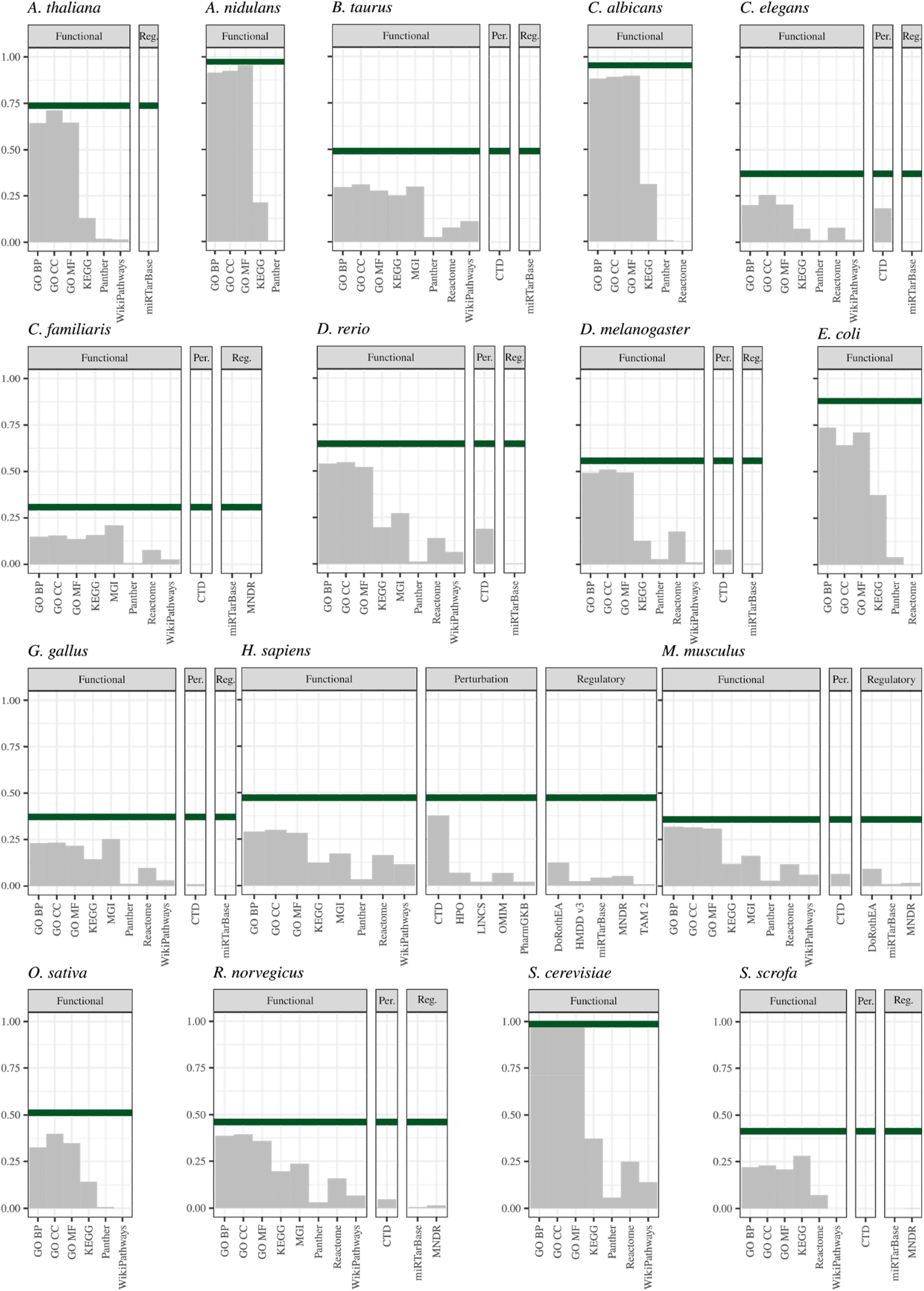
Gene coverage for each organism and annotation in the GeneCodis database. It is represented in a proportional scale, being 1 all the genes available in NCBI Entrez while the green line shows all the organism genes with at least one annotation from all the available. Annotations are divided in the three categories.

The index of genes and the inclusion of their different synonyms is built based on the NCBI gene info table (ftp://ftp.ncbi.nih.gov/gene/DATA/gene_info.gz) which includes Ensembl, Uniprot and HGCN Symbol identifiers. Additionally, Ensembl GTFs files are downloaded for each organism to expand and complete the Ensembl mapping with NCBI. miRNAs identifiers and accession numbers are mainly obtained from miRBase (Kozomara et al., 2019), however miRNAs symbols are also included in the mentioned NCBI table. CpG sites were annotated using human Infinium MethylationEPIC manifest file. Finally, TFs-gene annotations were retrieved from DoRothEA (Garcia-Alonso et al., 2019) which includes curated interactomes for human and mouse. DoRothEA provides different scoring TF-target interactions and those with the highest confidence (range A) were included.

#### Annotation data collection

GeneCodis 4 includes 19 different collections of annotations grouped into three main categories: functional, regulatory and perturbation annotations. Functional category cover the following databases: Gene Ontology (Gene Ontology Consortium, 2019) and its three subcategories, Biological Process, Molecular Function and Cellular Component, KEGG Pathways (Kanehisa et al., 2019), Mouse Genome Informatics database (Bult et al., 2019), Panther Pathways (Mi et al., 2021), Reactome (Jassal et al., 2020) and WikiPathways (Martens et al., 2021). Additionally, motivated by the necessity to assist global research on the SARS-CoV-2 virus, a GO subset which includes functions of human proteins used by the virus to enter a human cell is incorporated. The regulatory category contains two curated interactomes TF-gene pairs from DoRothEA and miRNA-gene interactions derived from strong evidence techniques such as: reporter assay, western blot and qRT-PCR from miRTarBase (Huang et al., 2020). Moreover, as a specialised regulatory subcategory, miRNAs-based functional annotations are added from the Tool for miRNA Set Analysis (TAM) database (Li et al., 2018), the Human microRNA Disease Database (HMDD) (Huang et al., 2019) and the Mammalian ncRNA-Disease Repository (MNDR) (Ning et al., 2021). miRNAs-based annotations mainly contain miRNAs-disease associations, only TAM includes functional categories but compared to gene-based databases it still lacks a comprehensive annotation, however it should be considered as the first option to overcome the miRNA enrichment analysis. Finally, perturbation category gathers two types of associations, 1) gene-chemicals that consist of the Comparative Toxicogenomics Database (CTD) (Davis et al., 2021), the LINCS consortium (Stathias et al., 2020) and PharmGKB (Thorn et al., 2013) databases and 2) gene-phenotype associations with the Human Phenotype Ontology (HPO) (Köhler et al., 2019) and the Online Mendelian Inheritance in Man (OMIM) (Rasmussen et al., 2020) database.

GeneCodis 4 annotation database can be downloaded from the help page for all the available organisms and gene nomenclatures. Additionally, the source URLs with the preprocessed files are provided.

### Web Tool Implementation

The GeneCodis 4 database is stored on a relational database built with PostgreSQL 12.

The GeneCodis 4 core algorithm is maintained from the previous version and is responsible for extracting coannotated terms in the form of closed frequent itemsets, particularly, all the possible combinations of terms that share a minimum number of input elements. Over this co- and single annotations, the statistical analysis is applied to find over-represented categories. Around this core, two Python3 libraries are employed to handle petitions and serve responses. GeneCodis 4 is developed as an application programming interface (API) using the Flask microframework which is then deployed via Gunicorn, a WSGI HTTP Server for UNIX. Finally, the GeneCodis 4 web page is sent to the client side via NGINX while also acting as a front-end reverse proxy of the application. GeneCodis 4 is deployed in Ubuntu Server 18, in a computer with 252GB of RAM memory and a microprocessor Intel(R) Xeon(R) Silver 4214R CPU @ 2.40GHz.

The GeneCodis 4 website is built with a simple integration of HTML, JavaScript and CSS. HTML pages are built with EJS, a templating language. Visualizations depend on two JavaScript libraries, D3.js and jQuery, plus DataTables plug-in.

### Analysis Workflow and Results Interpretation

GeneCodis 4 accepts different input lists types: genes, proteins, TFs, CpG sites or miRNAs. Although the input type must be specified, in contrast to other tools, the input is not limited to a single nomenclature nor need it to be specified.

It offers the option of studying a single list or two in a comparative analysis where the common and unique subsets are tested separately. By default, the application extracts annotations that are simultaneously present in at least 3 genes (min gene annotated), although this parameter can be modified by the user and is advisable to increment it when large lists are analysed. To optimize the balance among computational cost and processing time the co-annotation is limited to input lists of a maximum of 1000 elements and the simultaneous analysis of three different sources of annotations per analysis. The background set, also called universe, of genes, proteins, TFs, CpG sites or miRNAs matches the number of elements available in the gene data collection step. However, the user can introduce a custom input universe. Using a custom universe is recommendable, for example, if the user wants to test a list of genes and the upstream analysis is not based on NCBI annotation.

If the input list contains gene IDs, biological terms are directly retrieved from the annotation database. However, if the input list consists of TFs or CpG sites, the set of their target genes are defined before annotations can be linked using information from DoRothEA and Infinium MethylationEPIC annotation file, respectively. Finally, if miRNAs are provided as input list, terms can be directly linked using miRNAs-based annotations or miRNAs are transformed to their targets based on miRTarBase associations.

Once obtained the association input-terms, GeneCodis 4 computes a p-value based on the hypergeometric distribution and False Discovery Rate is used to correct for multiple testing. Enriched terms whose corrected p-value is not significant (Adj.Pval > 0.05) are discarded from the analysis. Furthermore, a relative enrichment score is computed as the proportion between two fractions: 1) the number of genes found in the input for an enriched annotation divided by the size of the input list; 2) all the genes under that annotation divided by all the genes in the database.

An interactive table is provided to explore the top 100 enriched terms and the entire table can be downloaded from the application website. Three interactive, customizable and downloadable visualizations are generated: a network where annotations are linked to their genes or miRNAs, a word cloud and a bar chart. The network has a profuse force layout which causes annotations to be close clustered as they share more genes. The size of the annotation node is proportional to −log10(Adj.Pval) and the color intensity goes in concordance with the number of genes found. The word cloud design varies, in co-annotation the terms appear according to the frequency that each single term is reported in the selected top co-annotated terms; meanwhile, in single analysis, the size of the word is related to its term −log10(Adj.Pval). Finally, in the bar chart, the length of each bar represents the −log10(Adj.Pval) and, likewise, the number of genes found are in consonance with the color intensity. Moreover, when miRNAs are enriched with gene-based annotations, along with the resulting visualizations, GeneCodis 4 provides a table that gathers the miRNAs targets interactions contemplated in our database.

## RESULTS

GeneCodis 4 database has been expanded including annotations of regulatory elements whose target interaction has a direct functional implication. It was accomplished by selecting miRNA-based annotations databases and two curated interactomes for TFs and miRNAs. Figure 1 offers a view of the coverage of GeneCodis 4 database for each organism and collection available.

GeneCodis 4 has been integrated with a third party application such as SRNAToolBox-sRNAbench (Aparicio-Puerta et al., 2019). By means of this integration, a complete analysis pipeline is presented. By using SRNAToolBox the user could perform sRNA expression profiling from next generation sequencing experiments data, in order to obtain a set of candidate miRNAs which can be, then, functionally characterised in GeneCodis 4 in a direct manner.

### Use Cases

#### Use case 1 - Gene-based annotations

In order to illustrate the new functionalities of GeneCodis 4, it were analyzed a set of most abundant miRNAs in exosomes secreted by EBV-transfected lymphoblastoids, a process that induces oncogenic-like properties including *in vitro* permanent growth (Koppers-Lalic et al., 2014). This data was obtained from Sequence Read Archive (SRA), run SRR1563060. Then, the miRNA profile was obtained using sRNAbench and subsequently sorted the result by Reads Per Million. The 5 most abundant human miRNAs were picked (hsa-miR-142-5p, hsa-miR-155-5p, hsa-miR-181a-5p, 4hsa-miR-21-5p, hsa-miR-92a-3p), all in its mature id, and removed Epstein-Barr virus miRNAs since this organism is not included in GeneCodis 4. From the sRNAbench results page, by clicking the GeneCodis 4 button included in the “More” section, the selected miRNAs were input in GeneCodis 4 and chose KEGG Pathways as annotation, the rest of the parameters were set to default.

The first part of the results is the input quality control, and it is noticeable that 5 miRNAs target 445 genes of which 299 are annotated in KEGG. Next, enrichment results are available in different tabs, the co-annotation and the single enrichment of KEGG. Not surprisingly, in both analyses, the term cancer is present in many of the KEGG pathways with significantly enriched terms, including “Pathways in cancer” and “MicroRNAs in cancer”. Figure 2 A) and B) show the top 12 terms obtained in the network and bar chart plot. Despite not including EBV miRNAs in the query, the terms “Epstein-Barr” or “herpes” (Epstein-Barr is a herpes virus) appears enriched in which indicates some sort of response to the infection. However, other terms like “Hepatitis B” also rank quite high even if they are far from the processes actually happening in this cell line. This can be attributed to the overlap between miRNAs normally secreted into exosomes and miRNAs that are abundant in the liver (Rekker et al., 2014; Shigoka et al., 2010). Although the use of indirect annotations can be quite useful to obtain some insight from miRNA data, this example goes to show the limitations of this approach as each miRNA target can be involved in many different pathways and significant enrichments can arise even from small lists of miRNAs.

**Figure 2.**
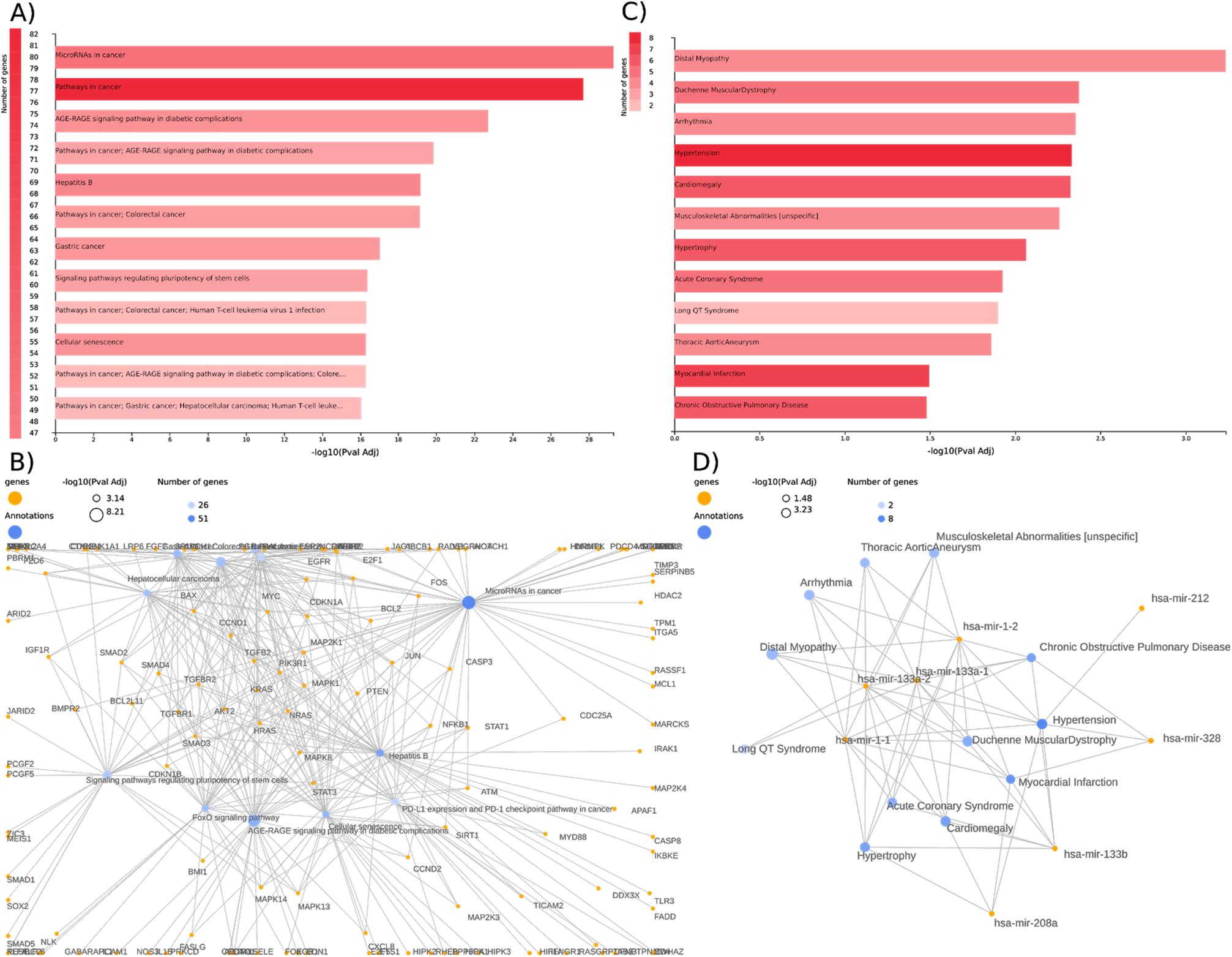
Bar chart and network plots obtained from GeneCodis 4. Representing A) co-annotation and B) singular KEGG enrichment from use case I; and C) barchart and D) network plot from singular HMDD enrichment in use case II.

#### Use case 2 - miRNA-based annotations

With the aim of exposing the aforementioned bias effect in miRNAs enrichment analyses and how GeneCodis 4 overcomes it, a set of miRNAs associated with a non-cancer-related pathology was chosen. These were searched for in the literature of heart related disorders, such as arrhythmia. The selected miRNAs (miR-1, miR-133, miR-328, miR-212, miR-208a) are extracted from the Table 1 in a review that studies miRNAs linked to cardiac excitability and other processes pertinent to arrhythmia (Kim, 2013). This list needs to be transformed to proper miRNAs identifiers because miR-1 and miR-133 refer to different families that share the same root sequence. Thus, following the latest miRBase annotation miR-1 family contains mir-1-1 and mir-1-2, likewise, miR-133 is divided into mir-133a-1, mir-133a-2 and mir-133b. Since the intention is to compare miRNAs-based and gene-based annotations, the list of miRNAs need to include their mature form, i.e. miR-133a-1, given that miRTarBase only associates mature forms with their target genes. Last, the human tag (hsa-) is incorporated to follow the miRBase identifiers convention. Gene-based annotations, Gene Ontology Biological Process, and miRNA-based, HMDD, are selected to test for enriched terms.

In this occasion, the list of 19 miRNAs leads to 231 targets, 13 miRNAs are annotated in HMDD and 137 targets and 2 miRNAs are annotated in KEGG. As in the previous use case, target genes cause mainly cancer-related KEGG terms being enriched. This is visible in single and co-annotation results. Notwithstanding, HMDD captures as the top significant terms, those mainly associated with muscle and heart disorders. The results from HMDD are plotted in Figure 2 where C) and D) shows the top 12 terms obtained in network and bar chart representations.

## DISCUSSION

GeneCodis 4 was released in February 2020 and, since then, it has received an average of 325 unique users per month, continuing with the previous trend that has positioned GeneCodis as a reference tool in the field of enrichment analysis. With this new release, GeneCodis have expanded the functionality of the application to cover data from methylation and miRNA profiling experiments. New and updated annotations and organisms along with a simplified interface aims to provide the users with a better experience. In relation with this, GeneCodis 4 incorporates exemplary input lists along with a small description, for different input types and organism selected to fit all types of experimental setting.

In conclusion, GeneCodis 4 provides an excellent web tool that requires no bioinformatics skills to integrate results from different omics granted its ability to analyse different regulatory elements and find concurrent annotations. The new network visualization permits customising the number of top enriched terms fulfilling a common need for the user, creating a compact plot to capture the most representative categories clustered and, at the same time, the number of genes that causes the enrichment.

This last update grants GeneCodis being still a singular type of enrichment tool thanks to its modular enrichment analysis whilst keeping up the latest demands in the biomedicine area.

## AVAILABILITY

GeneCodis 4 does not require login and is freely available at https://genecodis.genyo.es/.

## ACKNOWLEDGEMENT

We want to thank Manuel Orcera Herrera from GENyO’s IT management for his continuous technical support. Graphical abstract contains logos from referenced databases and icons made by Freepik, Smashicons and Eucalyp from www.flaticon.com.

## REFERENCES

Aparicio-Puerta, E., Lebrón, R., Rueda, A., Gómez-Martín, C., Giannoukakos, S., Jaspez, D., Medina, J.M., Zubkovic, A., Jurak, I., Fromm, B., et al. (2019). sRNAbench and sRNAtoolbox 2019: intuitive fast small RNA profiling and differential expression. Nucleic Acids Res 47, W530–W535.

Bleazard, T., Lamb, J.A., and Griffiths-Jones, S. (2015). Bias in microRNA functional enrichment analysis. Bioinformatics 31, 1592–1598.

Bult, C.J., Blake, J.A., Smith, C.L., Kadin, J.A., Richardson, J.E., and Mouse Genome Database Group (2019). Mouse Genome Database (MGD) 2019. Nucleic Acids Res 47, D801–D806.

Carmona-Saez, P., Chagoyen, M., Tirado, F., Carazo, J.M., and Pascual-Montano, A. (2007). GENECODIS: a web-based tool for finding significant concurrent annotations in gene lists. Genome Biol 8, R3.

Davis, A.P., Grondin, C.J., Johnson, R.J., Sciaky, D., Wiegers, J., Wiegers, T.C., and Mattingly, C.J. (2021). Comparative Toxicogenomics Database (CTD): update 2021. Nucleic Acids Res 49, D1138–D1143.

Garcia-Alonso, L., Holland, C.H., Ibrahim, M.M., Turei, D., and Saez-Rodriguez, J. (2019). Benchmark and integration of resources for the estimation of human transcription factor activities. Genome Res. 29, 1363–1375.

Garcia-Moreno, A., and Carmona-Saez, P. (2020). Computational Methods and Software Tools for Functional Analysis of miRNA Data. Biomolecules 10, 1252.

Geeleher, P., Hartnett, L., Egan, L.J., Golden, A., Raja Ali, R.A., and Seoighe, C. (2013). Gene-set analysis is severely biased when applied to genome-wide methylation data. Bioinformatics 29, 1851–1857.

Gene Ontology Consortium, T. (2019). The Gene Ontology Resource: 20 years and still GOing strong. Nucleic Acids Res 47, D330–D338.

Godard, P., and van Eyll, J. (2015). Pathway analysis from lists of microRNAs: common pitfalls and alternative strategy. Nucleic Acids Research 43, 3490–3497.

Huang, H.-Y., Lin, Y.-C.-D., Li, J., Huang, K.-Y., Shrestha, S., Hong, H.-C., Tang, Y., Chen, Y.-G., Jin, C.-N., Yu, Y., et al. (2020). miRTarBase 2020: updates to the experimentally validated microRNA-target interaction database. Nucleic Acids Res 48, D148–D154.

Huang, Z., Shi, J., Gao, Y., Cui, C., Zhang, S., Li, J., Zhou, Y., and Cui, Q. (2019). HMDD v3.0: a database for experimentally supported human microRNA-disease associations. Nucleic Acids Res 47, D1013–D1017.

Jassal, B., Matthews, L., Viteri, G., Gong, C., Lorente, P., Fabregat, A., Sidiropoulos, K., Cook, J., Gillespie, M., Haw, R., et al. (2020). The reactome pathway knowledgebase. Nucleic Acids Research 48, D498–D503.

Kanehisa, M., Sato, Y., Furumichi, M., Morishima, K., and Tanabe, M. (2019). New approach for understanding genome variations in KEGG. Nucleic Acids Res 47, D590–D595.

Kern, F., Fehlmann, T., Solomon, J., Schwed, L., Grammes, N., Backes, C., Van Keuren-Jensen, K., Craig, D.W., Meese, E., and Keller, A. (2020). miEAA 2.0: integrating multi-species microRNA enrichment analysis and workflow management systems. Nucleic Acids Res 48, W521–W528.

Kim, G.H. (2013). MicroRNA regulation of cardiac conduction and arrhythmias. Transl Res 161, 381–392.

Köhler, S., Carmody, L., Vasilevsky, N., Jacobsen, J.O.B., Danis, D., Gourdine, J.-P., Gargano, M., Harris, N.L., Matentzoglu, N., McMurry, J.A., et al. (2019). Expansion of the Human Phenotype Ontology (HPO) knowledge base and resources. Nucleic Acids Res 47, D1018–D1027.

Koppers-Lalic, D., Hackenberg, M., Bijnsdorp, I.V., van Eijndhoven, M.A.J., Sadek, P., Sie, D., Zini, N., Middeldorp, J.M., Ylstra, B., de Menezes, R.X., et al. (2014). Nontemplated nucleotide additions distinguish the small RNA composition in cells from exosomes. Cell Rep 8, 1649–1658.

Kozomara, A., Birgaoanu, M., and Griffiths-Jones, S. (2019). miRBase: from microRNA sequences to function. Nucleic Acids Res 47, D155–D162.

Kuleshov, M.V., Diaz, J.E.L., Flamholz, Z.N., Keenan, A.B., Lachmann, A., Wojciechowicz, M.L., Cagan, R.L., and Ma’ayan, A. (2019). modEnrichr: a suite of gene set enrichment analysis tools for model organisms. Nucleic Acids Res 47, W183–W190.

Li, J., Han, X., Wan, Y., Zhang, S., Zhao, Y., Fan, R., Cui, Q., and Zhou, Y. (2018). TAM 2.0: tool for MicroRNA set analysis. Nucleic Acids Res 46, W180–W185.

Martens, M., Ammar, A., Riutta, A., Waagmeester, A., Slenter, D.N., Hanspers, K., A Miller, R., Digles, D., Lopes, E.N., Ehrhart, F., et al. (2021). WikiPathways: connecting communities. Nucleic Acids Res 49, D613–D621.

McLean, C.Y., Bristor, D., Hiller, M., Clarke, S.L., Schaar, B.T., Lowe, C.B., Wenger, A.M., and Bejerano, G. (2010). GREAT improves functional interpretation of cis-regulatory regions. Nat Biotechnol 28, 495–501.

Mi, H., Ebert, D., Muruganujan, A., Mills, C., Albou, L.-P., Mushayamaha, T., and Thomas, P.D. (2021). PANTHER version 16: a revised family classification, tree-based classification tool, enhancer regions and extensive API. Nucleic Acids Res 49, D394–D403.

Ning, L., Cui, T., Zheng, B., Wang, N., Luo, J., Yang, B., Du, M., Cheng, J., Dou, Y., and Wang, D. (2021). MNDR v3.0: mammal ncRNA-disease repository with increased coverage and annotation. Nucleic Acids Res 49, D160–D164.

Nogales-Cadenas, R., Carmona-Saez, P., Vazquez, M., Vicente, C., Yang, X., Tirado, F., Carazo, J.M., and Pascual-Montano, A. (2009). GeneCodis: interpreting gene lists through enrichment analysis and integration of diverse biological information. Nucleic Acids Res 37, W317–322.

Paczkowska, M., Barenboim, J., Sintupisut, N., Fox, N.S., Zhu, H., Abd-Rabbo, D., Mee, M.W., Boutros, P.C., PCAWG Drivers and Functional Interpretation Working Group, Reimand, J., et al. (2020). Integrative pathway enrichment analysis of multivariate omics data. Nat Commun 11, 735.

Rasmussen, S.A., Hamosh, A., and OMIM curators (2020). What’s in a name? Issues to consider when naming Mendelian disorders. Genet Med 22, 1573–1575.

Rekker, K., Saare, M., Roost, A.M., Kubo, A.-L., Zarovni, N., Chiesi, A., Salumets, A., and Peters, M. (2014). Comparison of serum exosome isolation methods for microRNA profiling. Clin Biochem 47, 135–138.

Ren, X., and Kuan, P.F. (2019). methylGSA: a Bioconductor package and Shiny app for DNA methylation data length bias adjustment in gene set testing. Bioinformatics 35, 1958–1959.

Schübeler, D. (2015). Function and information content of DNA methylation. Nature 517, 321–326.

Shigoka, M., Tsuchida, A., Matsudo, T., Nagakawa, Y., Saito, H., Suzuki, Y., Aoki, T., Murakami, Y., Toyoda, H., Kumada, T., et al. (2010). Deregulation of miR-92a expression is implicated in hepatocellular carcinoma development. Pathol Int 60, 351–357.

Stathias, V., Turner, J., Koleti, A., Vidovic, D., Cooper, D., Fazel-Najafabadi, M., Pilarczyk, M., Terryn, R., Chung, C., Umeano, A., et al. (2020). LINCS Data Portal 2.0: next generation access point for perturbation-response signatures. Nucleic Acids Res 48, D431–D439.

Tabas-Madrid, D., Nogales-Cadenas, R., and Pascual-Montano, A. (2012). GeneCodis3: a non-redundant and modular enrichment analysis tool for functional genomics. Nucleic Acids Res 40, W478–483.

Thorn, C.F., Klein, T.E., and Altman, R.B. (2013). PharmGKB: the Pharmacogenomics Knowledge Base. Methods Mol Biol 1015, 311–320.

